# Explainable deep neural networks for predicting sample phenotypes from single-cell transcriptomics

**DOI:** 10.1101/2024.12.03.626549

**Authors:** Jordi Martorell-Marugán, Raúl López-Domínguez, Juan Antonio Villatoro-García, Daniel Toro-Domínguez, Marco Chierici, Giuseppe Jurman, Pedro Carmona-Sáez

## Abstract

Recent advances in single-cell RNA-Seq (scRNA-Seq) technologies have revolutionized our ability to gather molecular insights into different phenotypes, such as diseases, at the level of individual cells. The analysis of the resulting data poses significant challenges due to their sparsity and large volume, and proper statistical methods are required to analyze and extract information from scRNA-Seq datasets. Sample classification based on gene expression data has proven effective and valuable for precision medicine applications. However, standard classification schemas are often not suitable for scRNA-Seq due to its unique characteristics, and new algorithms are required to effectively analyze and classify samples at the single-cell level. In this article, we introduce singleDeep, an end-to-end pipeline that streamlines the analysis of scRNA-Seq data training deep neural networks, enabling robust prediction and characterization of sample phenotypes.

To validate the effectiveness of singleDeep, we applied it to make predictions on scRNA-Seq datasets from different conditions, including systemic lupus erythematosus and Alzheimer’s disease. Our results demonstrate strong diagnostic performance, validated both internally and externally. Moreover, compared with traditional machine learning methods applied to pseudobulk data, singleDeep consistently outperformed these approaches. In addition to prediction accuracy, singleDeep provides valuable insights into cell types and gene importance estimation for phenotypic characterization. This functionality provided additional and valuable information in our use cases. For instance, we corroborated that some interferon signature genes are consistently relevant for autoimmunity across all immune cell types in lupus. On the other hand, we discovered that genes linked to dementia have relevant roles in specific brain cell populations, such as APOE in astrocytes.

## Introduction

Transcriptomics experiments have been a fundamental tool to quantify gene expression within cells, offering a variety of applications unveiling numerous biological mechanisms and pathological insights across several diseases [1]. Traditional techniques, such as microarrays and bulk RNA-Seq, were founded on the assumption that gene expression within cells of the same tissue is uniform. Therefore, RNA from bulk populations of millions of cells is pooled and measured together, providing a generalized and oversmoothed gene expression profile for the entire tissue or sample. While these methods significantly advanced our understanding in different scientific domains [2], they overlooked inherent heterogeneity among cell types within tissues and the stochastic nature of gene expression (i.e., cell-to-cell differences in gene expression driven by randomness) [3]. As a result, addressing complex research questions, such as elucidating the roles of early developmental cells or understanding the functions of distinct cell types remained elusive using bulk approaches [4]. These limitations have been overcome by single-cell RNA-Sequencing (scRNA-Seq) technologies.

In scRNA-Seq experiments, the quantification of gene expression is extended to thousands of individual cells from each sample, offering an unprecedented depth of insight into cellular heterogeneity [5]. This approach has opened the door to groundbreaking discoveries [6], and cell atlases for different tissues and organisms are now available for the research community [7].

Analyzing scRNA-Seq data presents significant challenges due to its distinctive characteristics, especially high data volume and sparsity [8]. Consequently, standard methods for bulk transcriptomics data analysis are in most cases not suitable for scRNA-Seq. In the last few years, several methodologies have been developed to address scRNA-Seq data analysis challenges [9]. Well-established software packages such as the Seurat R toolkit [10] and the Scanpy Python library [11] offer functionalities for data processing, quality control, clustering, cell type identification or differential expression, among others. However, there is a lack of classification methods for predicting sample phenotypes from single-cell gene expression patterns.

Although machine learning (ML) methods have been successfully used for predicting clinical features such as disease status or treatment responses from bulk transcriptomics [12], they cannot be directly applied to scRNA-Seq data. Unlike bulk transcriptomics datasets, which typically provide a single value per gene and sample –the expected input for ML algorithms– scRNA-Seq datasets contain multiple values for each gene and sample, derived from individual cells. This introduces an additional layer of complexity that requires additional preprocessing to apply ML methods effectively. This can be partially addressed by generating pseudobulk data from scRNA-Seq by aggregating or averaging the expression counts across all cells within each sample [13]. However, this approach shows a major drawback as it cancels out the advantages of preserving information about the heterogeneity of gene expression within cell populations.

In this study, we introduce singleDeep, an innovative workflow grounded in deep learning (DL) to predict sample phenotypes from scRNA-Seq data. In detail, we implemented deep feed-forward artificial neural networks (ANNs) with an adaptative architecture that is adjusted automatically to the data. These ANNs are trained and tested for each cell population, taking gene expression from individual cells as input and the phenotype of their source samples as output.

Our methodology allowed us to attain precise predictions for complex phenotypes in genuine and practical contexts. In particular, we utilized scRNA-Seq data derived from blood samples to forecast the diagnosis of systemic lupus erythematosus (SLE), and from post-mortem brain samples to predict dementia associated to Alzheimer’s disease (AD). With a robust testing strategy, including validation with external data, singleDeep yielded good prediction performance in all cases, outperforming classical ML algorithms. Notably, in addition to the primary classification objective, we gained valuable biological insights from the singleDeep workflow, including the cell types that may be particularly relevant to the studied phenotype and the essential genes that contribute to the functional discrepancies among the sample groups within each cell population. We studied in detail the most relevant cell types and genes for each use case, being coherent with the previous knowledge of the diseases and revealing specific patterns at the cellular level.

## Results

### SingleDeep framework

SingleDeep has been implemented as a comprehensive framework encompassing all essential components for classification tasks in scRNA-Seq data (Figure 1). Available functionalities cover the following steps: **1) Data Preparation** compatible with standard Scanpy and Seurat objects. **2) Neural networks training** tailored to each cell population, ensuring specialized modeling for diverse cellular contexts. **3) Prediction and results generation**: singleDeep delivers robust predictions for test samples while also generating a detailed performance assessment of individual cell populations and the contributions of genes to the classification task. **4) Models export**, enabling their reuse for predicting phenotypes on new datasets. **5) External samples phenotyping** using previously trained neural networks to predict the phenotypes of new samples.

**Figure 1.**
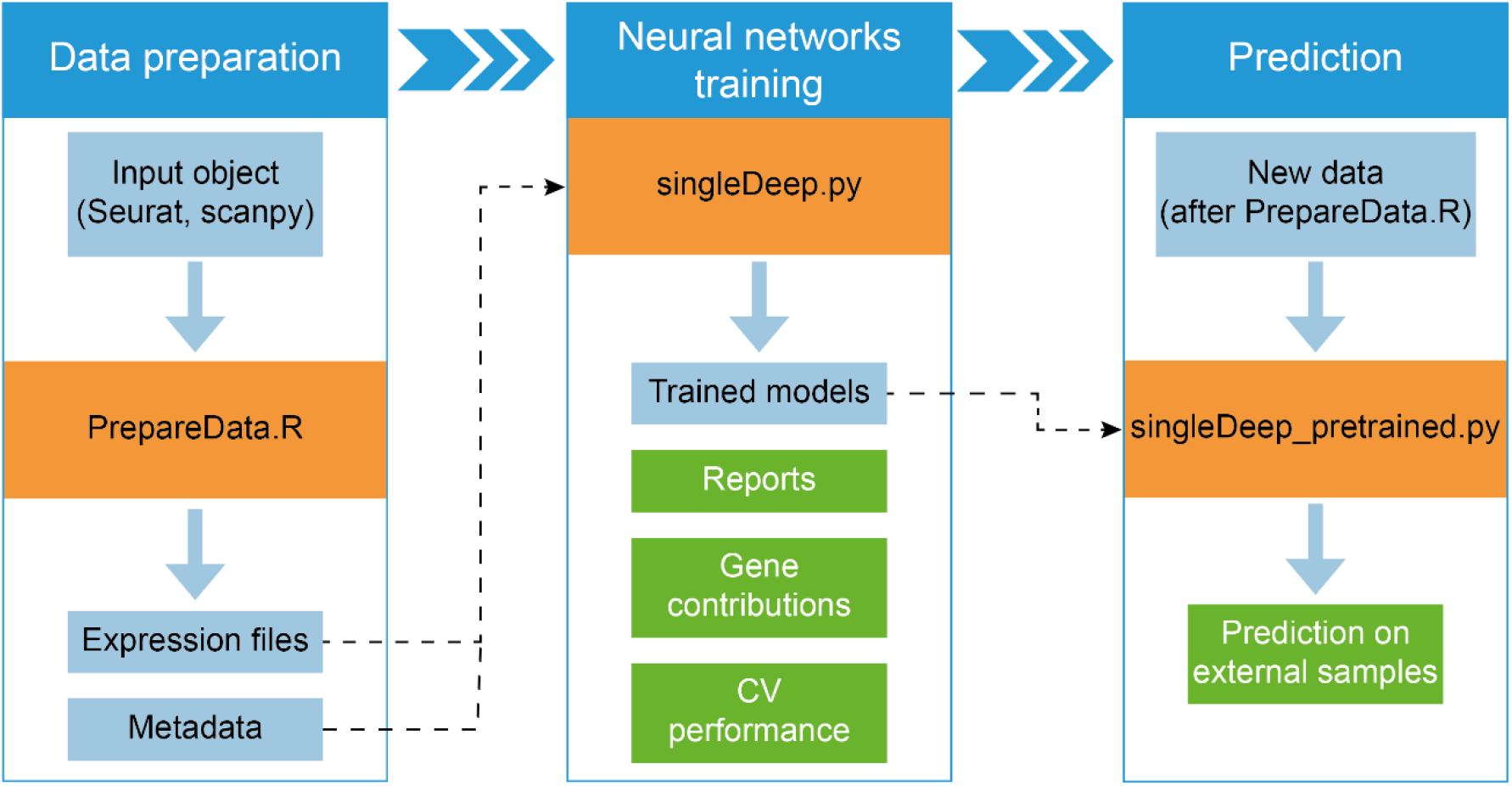
Scheme of the singleDeep pipeline. Blue boxes represent input and generated data, while orange boxes depict the scripts constituting the software. Green boxes represent the outputs. The workflow starts from standard scRNA-Seq objects processed with PrepareData.R to extract and store the expression matrix from each annotated cell population, along with its associated metadata and samples information. These files serve as input for singleDeep.py, which trains a dedicated neural network for each cell population and performs a nested cross-validation (CV) for internal validation. This step provides training reports, gene contributions and performance metrics. Lastly, singleDeep_pretrained.py is available to predict sample phenotypes on new external data using the trained networks from the previous step.

Each functionality incorporates numerous parameters that allow for fine-tuning of the analyses. Therefore, singleDeep offers a streamlined and adaptable solution for phenotype prediction. Software, detailed documentation and help pages for user guidance are publicly available at https://github.com/GENyO-BioInformatics/singleDeep.

### Lupus diagnosis is accurately predicted with singleDeep

To assess the effectiveness of singleDeep in a challenging clinical context, we conducted an extensive analysis of scRNA-Seq data obtained from a cohort comprising 162 SLE patients and 99 healthy controls (HCs) [14]. We validated the trained models with an external dataset, which included 40 pediatric and adult SLE patients and 16 healthy controls [15]. After training the ANNs and evaluating them using a stratified nested CV, singleDeep successfully predicted 228 out of 261 samples from the internal validation set, obtaining an accuracy of 87.36 % and a Matthew’s Correlation Coefficient (MCC) of 0.7297. Moreover, it predicts disease status of samples from the independent external test dataset with an accuracy of 83.93 % and a MCC of 0.8386. These results indicate that singleDeep was able to obtain high prediction metrics not only in the training data but also from external and independent data not used during the neural networks training.

We also compared our results with those obtained by applying some of the most widespread ML methods in pseudobulk gene expression data. SingleDeep consistently outperformed other approaches across all classification metrics during internal validation (Figure 2a) and demonstrated superior performance in most cases during external validation (Figure 2b).

**Figure 2.**
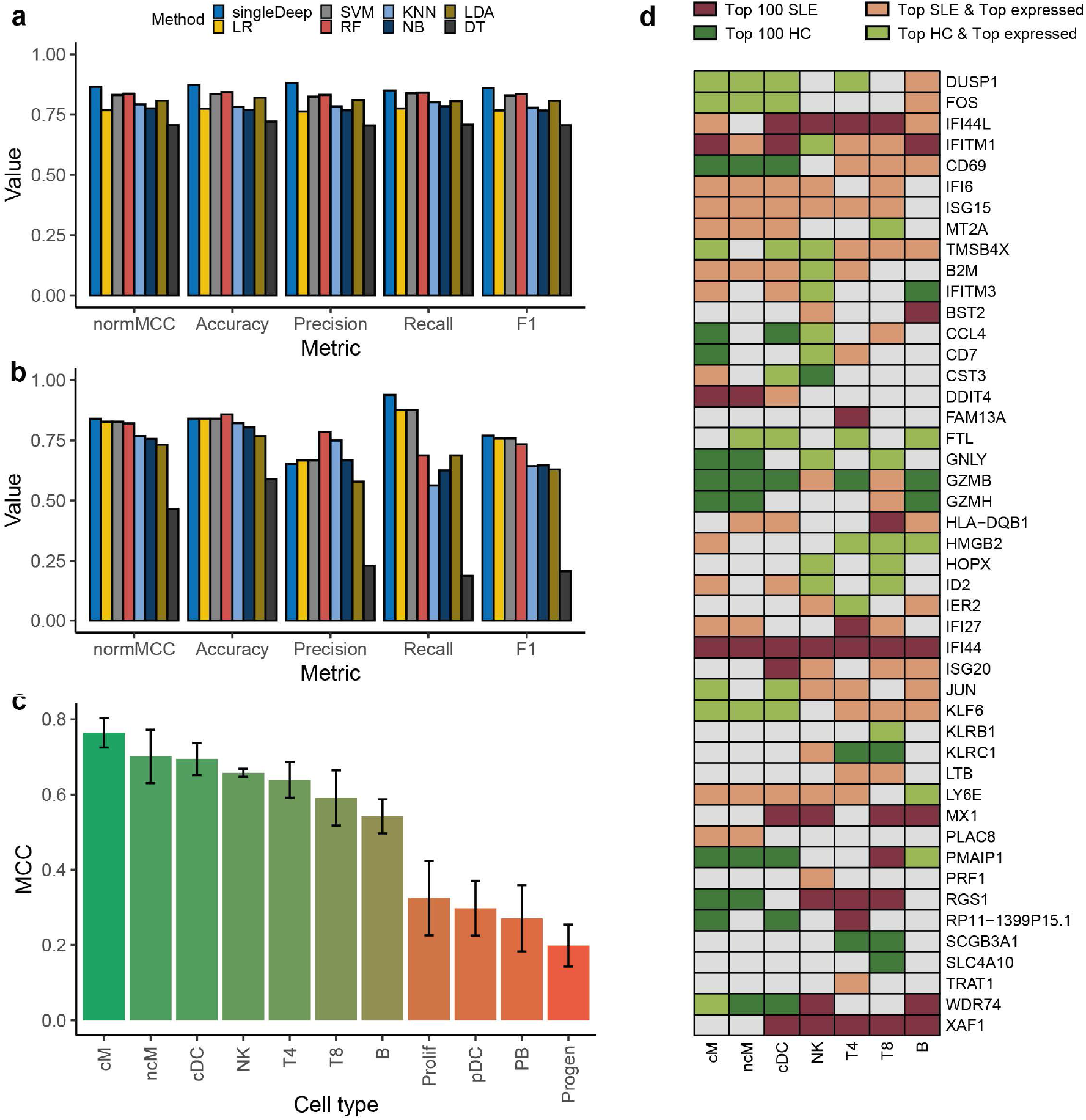
Assessment of SLE diagnosis prediction. **a** Comparative performance of singleDeep and classical ML algorithms on pseudobulk gene expression (LR: logistic regression, SVM: support vector machine, RF: random forest, KNN: K-nearest neighbors, NB: naïve Bayes, LDA: linear discriminant analysis, DT: decision trees). The metrics are computed for the test samples within the outer CV of the internal validation. **b** Performance metrics on the prediction of external data. **c** Mean MCC across cell types using the singleDeep workflow. Error bars denote standard deviations (SDs). **d** Key genes influencing sample classification as SLE or HC. These genes are among the top 10 contributors in at least one cell type. Colors indicate if genes are among the top 100 contributing to the classification of samples as SLE (red) or HC (green). Genes not within the top 100 for either category are depicted in gray cells. Lighter colors indicate genes that are among the top 5 % expressed genes in their cell type. Only cell types with a mean MCC > 0.4 are included.

Furthermore, singleDeep provides the functionality to evaluate the prediction performance for each specific cell population. We assessed the biological significance of these performances for SLE. Classical monocytes (cM) and nonclassical monocytes (ncM) emerged as the cell populations with the highest MCC values (Figure 2c), while others such as proliferating T and NK cells (Prolif), plasmacytoid dendritic cells (pDC), plasmablasts (PB) and progenitor cells (Progen) showed the lowest predictive performance. These findings are aligned with cell-type specific differential gene expression signature of SLE and control cases reported in the source study [14]. In addition, cMs were identified as the cells with the highest expression of the type I interferon-stimulated genes (ISGs) [14], one of the hallmarks of SLE. These results support the hypothesis that the predictive performance of individual cell populations provides valuable insights into the underlying mechanisms of the disease.

Next, we carried out a comprehensive assessment of gene contributions to the classification results across cell types (Supplementary Table 1). We visualized the top-contributing genes in a heatmap, highlighting top expressed genes in the corresponding cell types (Figure 2d). As can be observed, many of the top-contributing genes are highly expressed. It has been experimentally demonstrated that highly expressed genes are quantified erroneously in scRNA-Seq experiments [13]. Therefore, their importance for the classifications requires careful interpretation due to the possibility of being false positives.

Our findings reveal genes that consistently exhibit high relevance across multiple cell types, particularly ISGs such as IFI44L, IFI44, MX1 and XAF1 whose high expression is correlated to SLE[16]. We also found genes with a marked specificity for specific cell types, such as FAM13A in T CD4 (T4). Other genes are top contributors for classifying samples as SLE in some cell populations, while concurrently they are top contributors for classifying samples as HC in other cell populations (e.g., RGS1).

Remarkably, GZMH and GZMB emerge as top contributors to SLE classification in CD8 T cells (T8). The original work reported that a subpopulation of T8 cells overexpressing GZMH and GZMB was significantly enriched in SLE cases compared to controls (+8.6% and +6.0% for Asian and European populations respectively) [14]. Conversely, GZMK is one of the top contributors to the healthy phenotype in the same cell type, aligning with the previous study’s findings that a subpopulation of T8 cells with overexpressed GZMK is underrepresented in SLE patients (−1.1% and -0.7% for Asian and European populations respectively) [14]. Furthermore, our analysis indicates that CCR7 ranks as the 27^th^ contributor to healthy phenotypes in T4 cells (Supplementary Table 1). This may be due to the fact that naïve T4 cells expressing CCR7 were significantly underrepresented in SLE patients (Asian, –21.7%; European, –11.8%) [14]. These results suggest that the estimated contribution reflect differences between cell subpopulations abundance among phenotypes.

### Biological and technical factors influence the cell populations performance

We evaluated the impact of the influence of factors such as cell counts and the ratio of cells between classes in the predictive performance of individual cell types. Figure 3a shows the correlation of these measures with MCC values. We found that the ratio of cell abundances has no influence on the performance of the cell populations (Spearman’s correlation = -0.3, P-value = 0.3711). Conversely, a noticeable discrepancy in performance emerged between cell populations with a limited number of cells (< 10,000) and the remainder. Hence, there is significant positive correlation between the number of cells and MCC (Spearman’s correlation = 0.7782, P-value = 0.0048). These findings suggest that these factors contribute to the variations observed in prediction performance across cell types.

**Figure 3.**
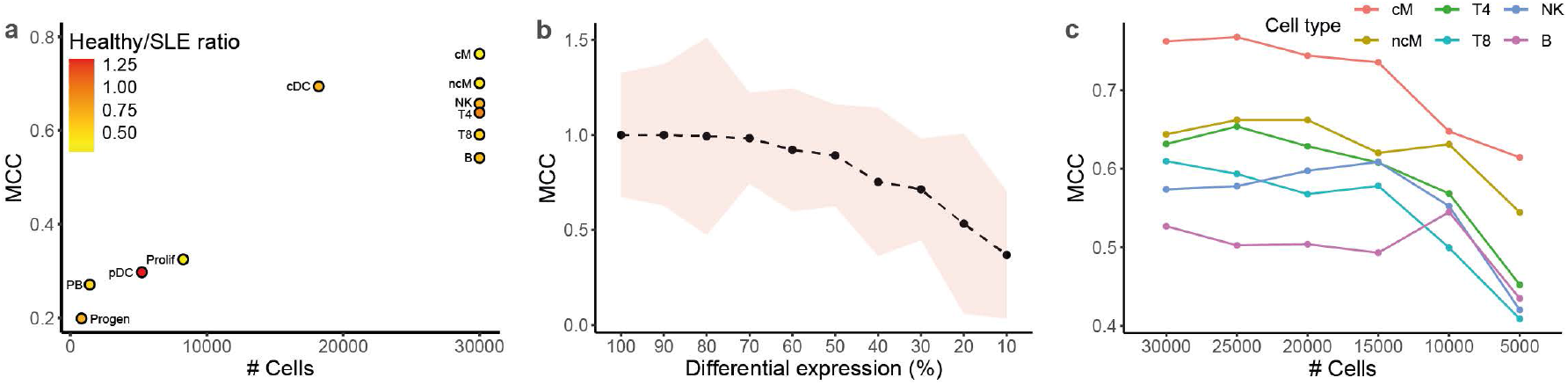
Evaluation of biological and technical factors affecting the cell types performance. **a** Relationship between the mean MCC and the number of cells used in the analysis for each cell type. Color indicates the ration between the number of cells originating from HC and SLE patients **b** Results from the simulation analysis. Each point represents the median MCC of a simulated cell population across 10 iterations. The shaded area represents the inter-quartile range. Differential expression levels for each cell population were systematically reduced to a certain percentage of their baseline (X-axis). **c** Ablation study for cell populations in the SLE dataset with > 30,000 cells. Each point represents the median MCC over 10 random undersampling iterations.

To further characterize this issue, we performed two complementary analyses: a simulation of synthetic data incorporating known differential expression levels between conditions and an ablation study using the SLE dataset. In the latter, we undersampled the cells progressively to assess the progressive impact on performance across cell types.

First, we simulated scRNA-Seq experiments for samples belonging to two conditions, establishing a baseline level of differential expression between them. This baseline differential expression varied across cell populations, ranging from no decrease (100% differential expression) to a reduction of 10% relative to the baseline. We used singleDeep to predict the condition of each sample and evaluate the performance of each cell population (Figure 3b). Our results demonstrate a gradual decline in cell population performance as the level of differential expression decreases to 80% of the baseline, demonstrating the faithful reflection of the level of differential expression between conditions by the MCC metric.

To further investigate the impact of cell number on the performance of cell types in our study, we conducted an ablation study focusing on populations containing more than 30,000 cells (i.e., cM, ncM, T4, T8, natural killer (NK), and B cells). Through random undersampling of cells, we repeated the analysis ten times for each cell count. Subsequently, each dataset underwent analysis with singleDeep, and the median MCC per cell type was computed at each ablation point (refer to Figure 3c). Notably, cM, ncM, and T4 consistently exhibited the best performance across varying cell counts, underscoring their high relevance in diagnosing SLE. However, other cell types displayed less consistent patterns, even demonstrating improved performance when analyzed with fewer cells (e.g., B cells performed optimally with 10,000 cells). Furthermore, direct comparisons between cell types with differing cell counts could potentially lead to misleading conclusions. For instance, if 30,000 sequenced T4 cells were compared to 5,000 ncM cells in an experiment, a direct MCC comparison might inaccurately suggest greater relevance of T4 cells in classification compared to ncM cells, without considering the discrepancy in cell numbers between the two populations.

In light of these findings, it becomes evident that both biological and technical factors contribute to the performance variations observed across cell types. Consequently, it is important to check the quality and characteristics of the data before drawing definitive conclusions from these results, considering the interplay between biological relevance and technical considerations.

### SingleDeep unravels cell-specific gene contributions to dementia

Finally, we aimed to evaluate singleDeep in solid tissues instead of blood samples, and in a context where transcriptional alterations are, a priori, more subtle than in an autoimmune disease or a viral infection. For that aim, we predicted dementia status in Alzheimer’s disease (AD) patients and controls from post-mortem brain dissections, specifically from the middle temporal gyrus (MTG) region. We analyzed the single-nucleus RNA-Seq (snRNA-Seq) dataset recently made available by the Seattle Alzheimer’s Disease Cell Atlas (SEA-AD) consortium [17] comprising 42 patients with dementia and 47 without dementia. We used the curated cell types annotations provided by the authors, who mapped their data to the recent reference single-cell brain atlas published by the BRAIN Initiative Cell Census Network (BICCN) [18]. The 1.378.211 cells were assigned to 18 reference cell types (Figure 4a) that were the input to singleDeep. An accuracy of 71.91% and a MCC of 0.4372 for the test samples of the nested CV was achieved (Supplementary Table 2). The MCC for individual cell types ranged from 0.1554 for cerebral cortex endothelial cells to 0.4476 for astrocytes of the cerebral cortex (Figure 4b and Supplementary Table 2).

**Figure 4.**
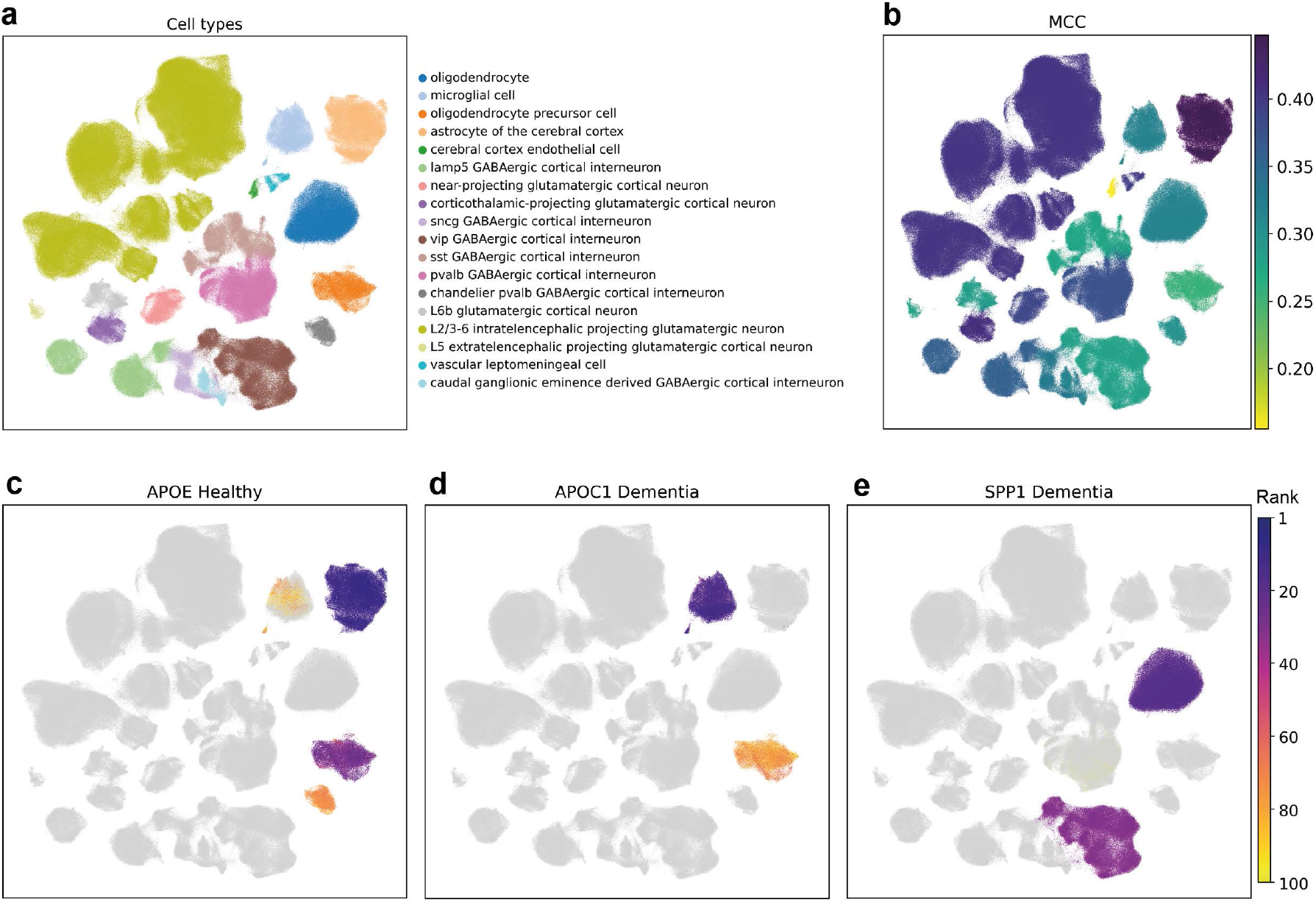
UMAP of the AD study cells colored by different criteria. **a** Cell types assigned by the authors of the source study. **b** Dementia prediction performance of singleDeep for individual cell types, represented by the MCC values. **c** APOE importance position to normal phenotype. Cells with APOE ranked under the 100^th^ position are colored in gray. **d** APOC1 importance position to AD phenotype. **e** SPP1 importance position to AD phenotype.

Several genes previously linked to AD were identified by singleDeep as top contributors to either dementia or normal states across different cell types (Supplementary Table 3). For instance, APOE is known for its strong genetic risk factor for AD, with one of its alleles (APOE4) increasing risk up to 15-fold [19]. In our study, APOE has been identified as top contributor to the healthy state in specific cell types, especially in astrocytes (Figure 4c). Interestingly, previous omics studies found downregulation of APOE4 in astrocytes compared to other APOE isoforms [20], possibly explaining its association with the classification of healthy donors in the singleDeep models.

On the other hand, the APOC1 gene, also associated with AD and whose expression affects cognitive functions [21], is a top contributor to dementia in microglial cells and to a lesser extent in oligodendrocyte precursor cells (Figure 4d). Moreover, the H2 allele of APOC1, linked to increased expression of this gene, is in genetic linkage disequilibrium with the APOE4 allele [22]. Consistent with our findings, previous snRNA-Seq studies in cortical samples have reported higher APOC1 expression in microglia of AD patients [23].

Furthermore, our analysis revealed a positive association between SPP1 gene expression and AD, particularly in oligodendrocytes and vip GABAergic cortical interneurons (Figure 4e). SPP1 is known to be implicated in neuroinflammation and various neurodegenerative diseases, including AD [24,25].

## Discussion

In this work we introduce singleDeep, a Python and R workflow for sample phenotype classification that uses deep learning with scRNA-Seq data. The workflow encompasses all essential analytical steps starting from the output of two widely used scRNA-Seq data processing software, Seurat and Scanpy.

Our results demonstrate that singleDeep achieves high performance in predicting disease status in three real cohorts under different scenarios. In the first place, we predicted the diagnosis of systemic lupus erythematosus (SLE) from blood samples. SLE is a complex autoimmune disease characterized by significant heterogeneity, making diagnosis challenging within clinical practice. Moreover, employing the trained models on an external independent SLE dataset yielded similarly good predictive performance. In both internal and external validations, our method outperformed in most of cases traditional machine learning (ML) algorithms applied to corresponding pseudobulk datasets.

Evaluation of prediction performance at cell type level by singleDeep underscores its relevance for the studied phenotype, although technical factors like the number of sequenced cells have also an impact on such performance. In our analysis, cM and ncM emerged as cell populations with the best classification scores, aligning with their known role in the type I interferon signature associated with SLE. Simulation and ablation analyses further support the relevance of cell types as predictive biomarkers of phenotypes, although technical factors must be considered before drawing conclusions.

The estimation of gene contributions by singleDeep provides relevant information, revealing different patterns. While certain interferon-stimulated genes (ISGs) consistently demonstrated high relevance across multiple cell types, indicating their association with SLE, others displayed cell type specificity or dual roles in sample classification.

In a second use case, we predicted dementia from brain post-mortem samples from Alzheimer’s disease patients and controls. We revealed that genes with major roles in dementia (i.e. APOE, APOC1 and SPP1) contribute to the predictions in specific cell types. These contributions are aligned to previous knowledge on their roles and differential expression in these cell types.

The singleDeep framework represents a significant advancement in the analysis of scRNA-Seq data, providing a comprehensive solution for phenotype prediction and biological interpretation. Furthermore, the adaptability of the singleDeep framework to adjust the neural networks design represents a notable strength. The ANN design (e.g., number of layers) may be modified, as well as the architecture itself (e.g. convolutional neural networks could be used instead of feed-forward neural networks).

Some previous methods exist for phenotypes prediction using scRNA-Seq data [26–29]. However, these methods lack the capability to export trained models and predict phenotypes on external datasets. Therefore, we could not compare singleDeep to these methods with an external validation strategy, which is crucial for assessing the generalizability of the trained models. Additionally, these methods do not systematically evaluate the importance of genes for predictions across different cell types. In our use cases, we demonstrated that this assessment provides biologically insightful information, revealing which genes are most relevant in each cell type for the studied phenotype.

## Materials and methods

### SingleDeep workflow

The starting point of singleDeep is a processed scRNA-Seq dataset following standard preprocessing steps saved as either a Seurat [10] or Scanpy [11] object. Optionally, users can specify a maximum number of cells per cell population to reduce the computational cost of the analysis. If this number is exceeded, the populations are randomly undersampled to the specified number.

Subsequently, a nested CV is employed to train and test ANNs. Each inner and outer fold involves splitting samples into a training set for network training and a test set for assessing network efficacy. Optionally, gene-wise standard scaling is performed using the mean and SD from the training data. A modified version of the Scanpy’s scale function [11] is used for this purpose.

For each cell population, the parameters of the ANN and the MCC [30,31] are calculated using all cells from the training samples. MCC, which is specially suitable for class-imbalanced data [32,33] is calculated for binary classifications as:

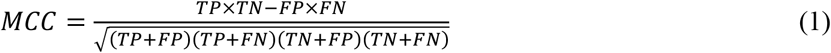

while for multiclass classifications is computed as:

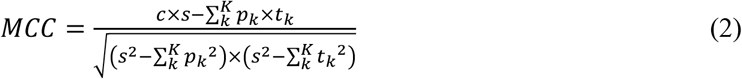

Where TP is the number of true positives, TN the number of true negatives, FP the number of false positives and FN the number of false negatives, t_k_ the number of *k* labelled samples in the data, p_k_ the number of times that class *k* was predicted; c the number of samples correctly predicted and s the total number of samples. The MCC values range from 1 (perfect prediction) to -1 (perfect opposite prediction)[30].

Once the phenotypes of the individual cells of test samples are predicted, the overall sample phenotype is assigned based on the majority prediction within that cell population. This process is repeated independently for each cell population, resulting in multiple predictions per sample. A weighted voting process is computed to assign final sample classes. The weights of the cell populations (W_c_) determined by their performance during the inner CV loop are calculated as:

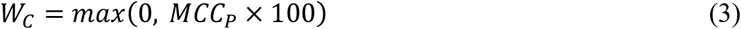

Being MCC_P_ the MCC of the test samples for cell population P.

SingleDeep refines a final model for each cell type using the entire dataset for two purposes: estimating gene contributions to the classification and saving the model for external data use. This final model is trained with the entire dataset, using the average optimal number of epochs from the inner CV. SingleDeep yields output files encapsulating performance metrics of overall and individual cell type predictions, along with gene contributions, both delineated by CV fold and the aggregate across all folds. Additionally, TensorBoard[34] reports are generated for each iteration and cell population to generate visualizations of training and test processes. Figure 5 provides a schematic representation of the singleDeep workflow and includes examples of predictions for a simple scenario.

**Figure 5.**
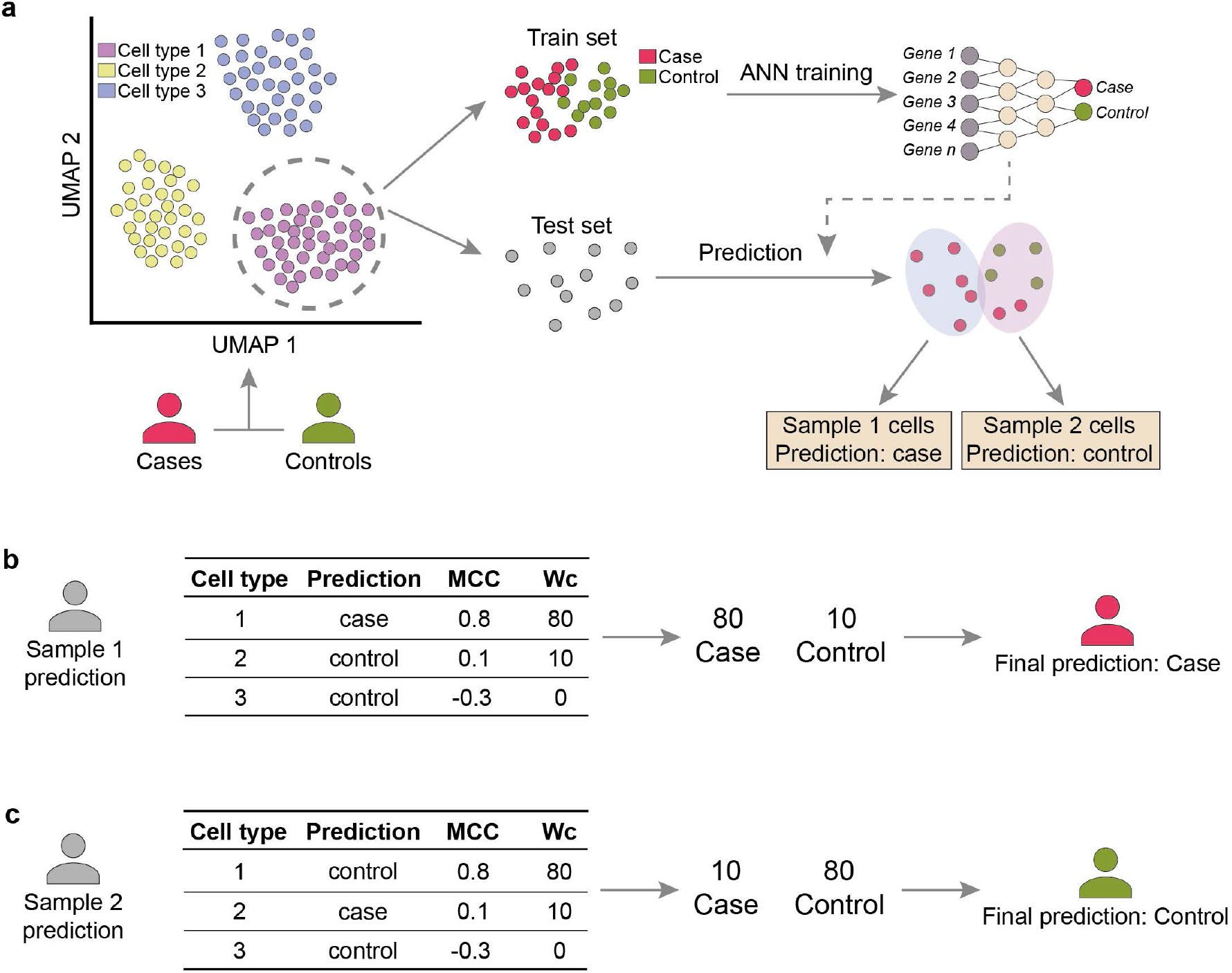
SingleDeep workflow in a simplified scenario. **a** Starting from a processed scRNA-Seq dataset comprising samples with two phenotypes (case and control), cells from each cell population are split into train and test sets. Train sets are used for training the ANN, which is used to predict phenotypes for the test sets. Predictions for test samples are determined by the most common phenotype predicted for their cells. This process is independently performed for each cell population. **b** Final prediction example for a case sample. The table contains predictions derived from individual cell populations, along with associated MCC values and weights. The final prediction is based on a weighted voting. **c** Analogous to b, presenting a prediction example for a control sample.

### Neural network architecture and training

The singleDeep framework leverages the PyTorch library [35] to build and train DL models. The default architecture consists of a 5-layer fully connected feed-forward neural network, with an adaptative number of nodes in each layer. The input layer includes as many nodes as the number of genes in the input dataset. Subsequent hidden layers progressively reduce in node count: the first hidden layer has half the nodes of the input layer, the second hidden layer has half of the first, and the third has a quarter of the second hidden. The output layer contains *k* nodes, being *k* the number of classes in the classification problem. Rectified linear Units (ReLU) [36] are used as activation functions in the hidden layers, and are calculated as:

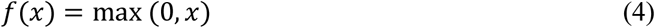

Where *x* is the intput to the node.

Training involves mini-batch processing, with the batch size dynamically set as a percentage of the training cells (defaulting to 10%).

The optimization process employs stochastic gradient descent with an adjustable learning rate to minimize the loss function. As loss function, the cross-entropy loss (L_CE_) for *k* classes (equation (5)) is used.

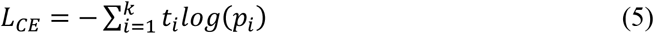

Where t_i_ is the real label and p_i_ the probability for the *i*^*th*^ class, calculated with the Softmax function [37]. Weights inversely proportional to the proportion of cells from each class are introduced to deal with potential imbalanced data.

The number of training epochs can be customized by the user. Nonetheless, to prevent overfitting on training data, an early stopping mechanism based on the loss function of the test set is implemented. This mechanism involves two sliding windows of 5 epochs each, separated by 2 epochs. If the difference in mean loss between these windows surpasses a threshold, training is stopped. We implemented this windows-based early stopping to avoid that a punctual increase in the testing loss stops prematurely the training. Before early stopping, a minimum of training epochs must have been performed to avoid premature stopping. Model parameters are saved at each new minimum loss, ensuring recovery of the model with the optimal performance. For training with the entire dataset, the number of epochs is set equal to the mean optimal number derived from the outer CV loop.

### Gene contributions estimation

Gene contributions to the predictions are estimated with the DeepLIFT [38] approach, implemented in the Captum library [39]. DeepLIFT is a back-propagation-based strategy for DL models interpretability that use the differences between the input values and a baseline, as well as the difference between the predictions for a target class using the baseline and the input values. To mitigate the bias towards highly expressed genes, we use local contributions, which do not multiply the contributions by the difference between the input expression and the baseline, as opposed to the predefined global contributions.

In singleDeep, the minimum expression of each gene serves as the baseline to make the contributions easily interpretable. Positive contributions indicate that a gene’s expression contributes to the cell being classified as the target class, while negative contributions indicate the opposite. This target class is chosen by the user.

To make the contributions comparable across samples, they are computed on the final model trained with all data. Gene contributions are averaged to get a unique value for each sample and cell population.

### Use cases data acquisition and processing

The training and validation datasets for SLE were retrieved from the NCBI GEO [40] repository (codes GSE174188 and GSE135779 respectively). We preprocessed the raw gene counts likewise the training data source study [14] using Scanpy [11]. This involved gene and cell filtering, batch effect correction, regression by total and mitochondrial counts, and gene expression normalization. The top 2000 highly variable genes were identified from the training dataset, and both datasets were subsequently sampled to include only these genes. Independent scaling of gene expression was performed for each dataset. To ensure uniformity in cell type annotation, we utilized Scanpy’s ingest method to project cell types from the validation dataset into the training dataset, adopting the cell type annotation provided by the authors of the source study [14]. For the training and internal validation with singleDeep, a nested CV with 5 outer folds and 4 inner folds was used.

The AD dataset was retrieved from the Chan Zuckerberg CELLxGENE Explorer [41]. We selected the top 2000 highly variable genes and we scaled the data. We used 3 outer and inner folds in the nested CVs.

### Performance metrics

In addition to MCC, singleDeep returns additional performance metrics: Accuracy (equation (6)), Precision (equation (7)), Recall (equation (8)) and F1 score (equation (9)).

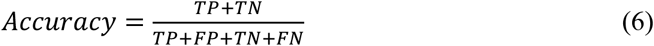

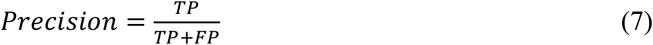

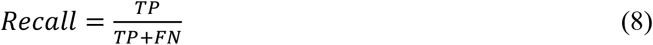

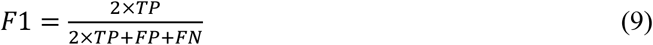

Being TP the number of true positives, FP the number of false positives, TN the number of true negatives and FN the number of false negatives.

Furthermore, to make the MCC comparable to the rest of the metrics, we calculated the normMCC [33] following equation (10).

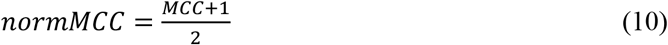

### Classification with traditional ML algorithms

In assessing the performance of singleDeep against traditional machine learning (ML) algorithms, we generated pseudobulk RNA-Seq datasets for both the training and validation datasets. For that aim, we calculated the mean normalized expression across all cells within each sample. We used the scikit-learn Python library [42] for training and validating models using LR, SVM, RF, KNN, NB, LDA and DT algorithms. In parallel to the singleDeep analysis, we used a 5×4 nested CV, using the inner CV for hyperparameters optimization and the outer CV for internal validation. For each method, we performed 100 iterations involving a random combination of hyperparameter values. Within each outer fold, we used the best combination of hyperparameters determined through the inner CV to train a model with the training samples and predict labels for the test samples. To generate the final model for external validation, we retrained the model with the entire dataset, using the hyperparameters from the outer CV fold with the highest MCC.

### Synthetic data generation

To investigate the impact of known differential expression on individual cell types, we employed the splatter R package [43,44] to generate simulated datasets. We generated scRNA-Seq experiments for 100 samples belonging to two distinct conditions. Each sample comprised 10 cell populations, each consisting of 100 cells expressing 2000 genes.

To introduce differential expression, we established a baseline of 5% variation between conditions for 50 genes. We customized the splatter functions to reduce this baseline differential expression increasingly across cell populations, spanning from no reduction (100% differential expression) to a 10% difference relative to the baseline. Additionally, we introduced a custom parameter to add extra variability between samples by nullifying the differential expression of 10% randomly selected genes per sample.

To account for the inherent randomness introduced during data simulation, we executed ten simulation iterations, calculating the median MCC for each cell type across these iterations.

## Data, Materials, and Software Availability

Synthetic datasets used in this study were generated using the splatter R package with custom modifications and may be generated using our shared code. The SLE datasets analyzed during the current study were generated by Perez et al. [14] and Nehar-Belaid et al. [15] and are available in the NCBI GEO repository (accession codes GSE174188 and GSE135779, respectively. The AD dataset was generated by the SEA-AD consortium [17] and is available in the Chan Zuckerberg CELLxCELL Explorer.

Source code for the singleDeep library is available on GitHub (https://github.com/GENyO-BioInformatics/singleDeep). Code used for the analyses and figures of this study is available at https://github.com/GENyO-BioInformatics/singleDeep_article.

## Acknowledgements

This work was funded by grant PID2020-119032RB-I00 funded by MCIN/AEI/10.13039/501100011033 and FEDER/Junta de Andalucía-Consejería de Universidad, Investigación e Innovación (ProyExcel_00978, P20_00335, B‐CTS‐40‐UGR20). RLD is supported by “Ayudas para personal técnico de apoyo”, PTA2021-021013-I from Agencia estatal de Investigación – Ministerio de Ciencia e Innovación. The authors thank the Supercomputing and Bioinnovation Center (SCBI) of the University of Malaga for their provision of computational resources (the supercomputer Picasso) and technical support (www.scbi.uma.es/site).

## Author contributions

PCS: Conceptualization (lead), supervision (lead), methodology (supporting), writing – original draft (lead). JMM: Software (lead), formal analysis (lead), methodology (lead), writing – original draft (lead). RLD, JAVG and DTD: formal analysis (supporting). MC and GJ: Software (supporting). All authors contributed to writing, review and editing the final manuscript.

## Key points

- singleDeep framework includes all the steps for samples phenotype prediction with scRNA-Seq data
- Cell types and genes importance for classification provide relevant biological information
- Complex diseases’ heterogeneity may be unveiled with singleDeep and single-cell transcriptomics

## Descriptions of the authors

**Jordi Martorell-Marugan** is a postdoctoral researcher in Bioinformatics and omics data analysis in biomedicine. He is interested in the development of new omics data analysis algorithms, especially based on machine learning and systems biology.

**Raul López-Domínguez** is a biologist and bioinformatician with expertise in Data Science applied to immunology.

**Juan Antonio Villatoro-García** is a statistician specialized in Biostatistics and data integration from different sources.

**Daniel Toro-Domínguez** is a postdoctoral researcher in Karolinska Institutet and is expert in Computational Biology applied to the study of autoimmune diseases, omics data analysis, machine learning and molecular clustering analysis.

**Marco Chierici** is a senior researcher, with a PhD in Bioengineering, at the Data Science for Health (DSH) Research Unit. His main research interests include the integration of artificial intelligence in computational biology frameworks. He is an expert on bioinformatics tools for massive omics, high-performance computing, and scientific programming in Python and R.

**Giuseppe Jurman** is a mathematician, with a PhD in Algebra, Head of the Data Science for Health (DSH) Research Unit, working on various aspects of data science, especially for computational biology.

**Pedro Carmona-Sáez** is lecturer in the Statistics and Operational Research department of the University of Granada (Granada, Spain). He mainly engages in bioinformatics research, focusing on developing methods for integrating omics data to decipher molecular mechanisms of complex diseases.

