## Supplementary Table 2 for "Explainable deep neural networks for predicting sample phenotypes from single-cell transcriptomics"

**Supplementary Table 2.** Performance of the overall model and the individual cell types for the prediction of dementia. Metrics for the individual cell types correspond to the average of the external folds of the nested CV.

| **Cell type** | **Accuracy** | **Precision** | **Recall** | **F1** | **MCC** |
| --- | --- | --- | --- | --- | --- |
| Overall | 0.7191 | 0.7391 | 0.7234 | 0.7312 | 0.4372 |
| astrocyte of the cerebral cortex | 0.7195 | 0.7591 | 0.6935 | 0.7206 | 0.4475 |
| corticothalamic-projecting glutamatergic cortical neuron | 0.7008 | 0.7493 | 0.6578 | 0.6985 | 0.4095 |
| L2_3-6 intratelencephalic projecting glutamatergic neuron | 0.7008 | 0.7285 | 0.6961 | 0.7112 | 0.4025 |
| vascular leptomeningeal cell | 0.6927 | 0.7216 | 0.7083 | 0.7067 | 0.3950 |
| near-projecting glutamatergic cortical neuron | 0.6897 | 0.7296 | 0.6649 | 0.6927 | 0.3853 |
| pvalb GABAergic cortical interneuron | 0.6878 | 0.7110 | 0.6961 | 0.7024 | 0.3755 |
| lamp5 GABAergic cortical interneuron | 0.6748 | 0.7219 | 0.6336 | 0.6701 | 0.3595 |
| sncg GABAergic cortical interneuron | 0.6533 | 0.7326 | 0.5388 | 0.6181 | 0.3293 |
| microglial cell | 0.6507 | 0.7156 | 0.5707 | 0.6318 | 0.3177 |
| oligodendrocyte | 0.6438 | 0.7322 | 0.5177 | 0.5990 | 0.3160 |
| caudal ganglionic eminence derived GABAergic cortical interneuron | 0.6489 | 0.7158 | 0.5603 | 0.6244 | 0.3157 |
| chandelier pvalb GABAergic cortical interneuron | 0.6528 | 0.6665 | 0.7383 | 0.6960 | 0.3023 |
| L6b glutamatergic cortical neuron | 0.6223 | 0.7382 | 0.4727 | 0.5732 | 0.2868 |
| vip GABAergic cortical interneuron | 0.6358 | 0.6764 | 0.6128 | 0.6339 | 0.2813 |
| L5 extratelencephalic projecting glutamatergic cortical neuron | 0.6274 | 0.7133 | 0.5253 | 0.5958 | 0.2804 |
| sst GABAergic cortical interneuron | 0.6135 | 0.7433 | 0.4344 | 0.5309 | 0.2766 |
| oligodendrocyte precursor cell | 0.6227 | 0.6769 | 0.5561 | 0.6103 | 0.2562 |
| cerebral cortex endothelial cell | 0.5650 | 0.6387 | 0.4732 | 0.5255 | 0.1554 |
